# Opening Pandora’s box: high level resistance to antibiotics of last resort in Gram negative bacteria from Nigeria

**DOI:** 10.1101/344093

**Authors:** David O Ogbolu, Laura J V Piddock, Mark A Webber

## Abstract

Antimicrobial resistance (AMR) is a global problem but information about the prevalence and mechanisms of resistance in sub-Saharan Africa are lacking. We determined the percentage of drug resistant isolates and resistance mechanisms in 307 Gram negative isolates randomly collected from south western Nigeria. Susceptibility testing revealed 78.1%, 92.2% and 52.6% of all isolates were resistant to fluoroquinolones, third generation cephalosporins and carbapenems respectively. There were more resistant isolates from the stools of uninfected patients than from specimens of patients with symptoms of infections. Only a small proportion of *E. coli* (10%) and *Klebsiella* (7%) isolates produced a carbapenemase. Whole genome sequencing of selected isolates identified the presence of globally disseminated clones. This depicts a crisis for the use of first line therapy in Nigerian patients, it is likely that Nigeria is playing a significant role in the spread of AMR due to her high population and mobility across the globe.

## INTRODUCTION

Nigeria and other sub-Saharan African countries face an increasing number of healthcare associated infections caused by multi-drug resistant (MDR) Gram-negative bacteria[1, 2]. Pathogenic species have evolved resistance to multiple antimicrobial agents including the mainstays of treatment[3, 4]. This is of particular concern as there are few new antibiotics in development with activity against Gram-negative bacteria[5]. Whilst Gram-negative bacteria are often intrinsically more resistant to many antibiotics than Gram-positive species, drug resistance to the most clinically important antibiotics is largely mediated by genes which are transmitted on plasmids that can readily spread through bacterial populations[6]. Species belonging to the *Enterobacteriaceae* family are the most commonly isolated MDR bacteria causing healthcare associated infections globally including in sub-Saharan Africa[7]. These bacteria include extended-spectrum β-lactamase (ESBL) producing *K. pneumoniae* and *E. coli* which are associated with both hospital and community infections with very high mortality rates[8. 9].

Carbapenems have become a mainstay of therapy for the treatment of ESBL producing Gram-negative bacteria. This has led to an increase in carbapenem use for treatment of serious infections[10]. As a result, there has been a selective pressure for carbapenem resistance and carbapenem resistant strains have spread globally [11](11). Worryingly, carbapenem-resistant bacteria are often resistant to other classes of antibiotics including aminoglycosides, fluoroquinolones and other ß-lactams, with often colistin and tigecycline as the only effective drugs. Resistance to both these antibiotics can also easily evolve making them unreliable as ‘last resort’ therapies[12, 13].

Mechanisms of carbapenem resistance include production of carbapenemase enzymes and/or repression of porins to limit entry of these drugs into the bacterial cell. Carbapenemases belong to different enzyme families including the metallo-carbapenemases (including the NDM and VIM enzymes) or non-metallo-types (including the KPC and OXA-48 enzymes)[14]. Some of these enzymes are associated with a particular species, for example the ‘*Klebsiella Pneumoniae* Carbapenemase’ or ‘KPC’ enzymes are typically found in *K. pneumoniae* species[15]. Other enzymes, such as NDM are found in many species [16].

Nigeria is often referred to as the “Giant of Africa”, owing to its large population and economy with approximately 182 million inhabitants, by far the largest in Africa. The Nigerian population is highly mobile and over 70% of Nigerians are under the age of 50. The large size and high level of mobility of this population makes import and export of antibiotic resistant bacteria a real concern for both Nigeria, but also the wider global community. Gram-negative bacteria cause a significant number of infections in Nigerian hospitals and represent the majority of isolates from both wounds and urinary tract infections; these form the largest group of clinical specimens received in Nigerian clinical microbiology laboratories[1]. Carbapenem-resistant Gram-negative bacteria have become prevalent in many parts of the world including Nigeria and sub Saharan Africa. However, to our knowledge there are few data and no organized antimicrobial resistance (AMR) surveillance networks for Africa. Recently, we showed that carbapenem resistant bacteria are present in Nigeria. The details of the strains with this phenotype and the mechanisms of resistance have not been studied in detail [3, 4](3,4). The potential role of the Nigerian population in the global spread of antibiotic resistance is great but the local situation is not understood. We determined a retrospective analysis of percentage resistance and resistance mechanisms in clinical and commensal isolates of Gram negative bacteria.

## METHODS

### Sample sites and bacterial isolates

The majority of the Nigerian population is found in the southwest of the country; this is also where major transportation hubs are located. Gram negative bacterial isolates for this study were recovered from patients admitted to three large teaching hospitals located in three states of south western Nigeria from a range of clinical specimens with invasive and colonized infections (Table 1, and Figure 1). The Olabisi Onabanjo University Teaching Hospital (OOUTH) is a 185-bed tertiary health care center and major referral center for Ogun State. The University College hospital (UCH) is in Ibadan in Oyo state and has 850 beds. The Obafemi Awolowo University Teaching Hospitals Complex (OAUTHC) is a teaching hospital with 535 beds and is in Osun state.

**Table 1.**
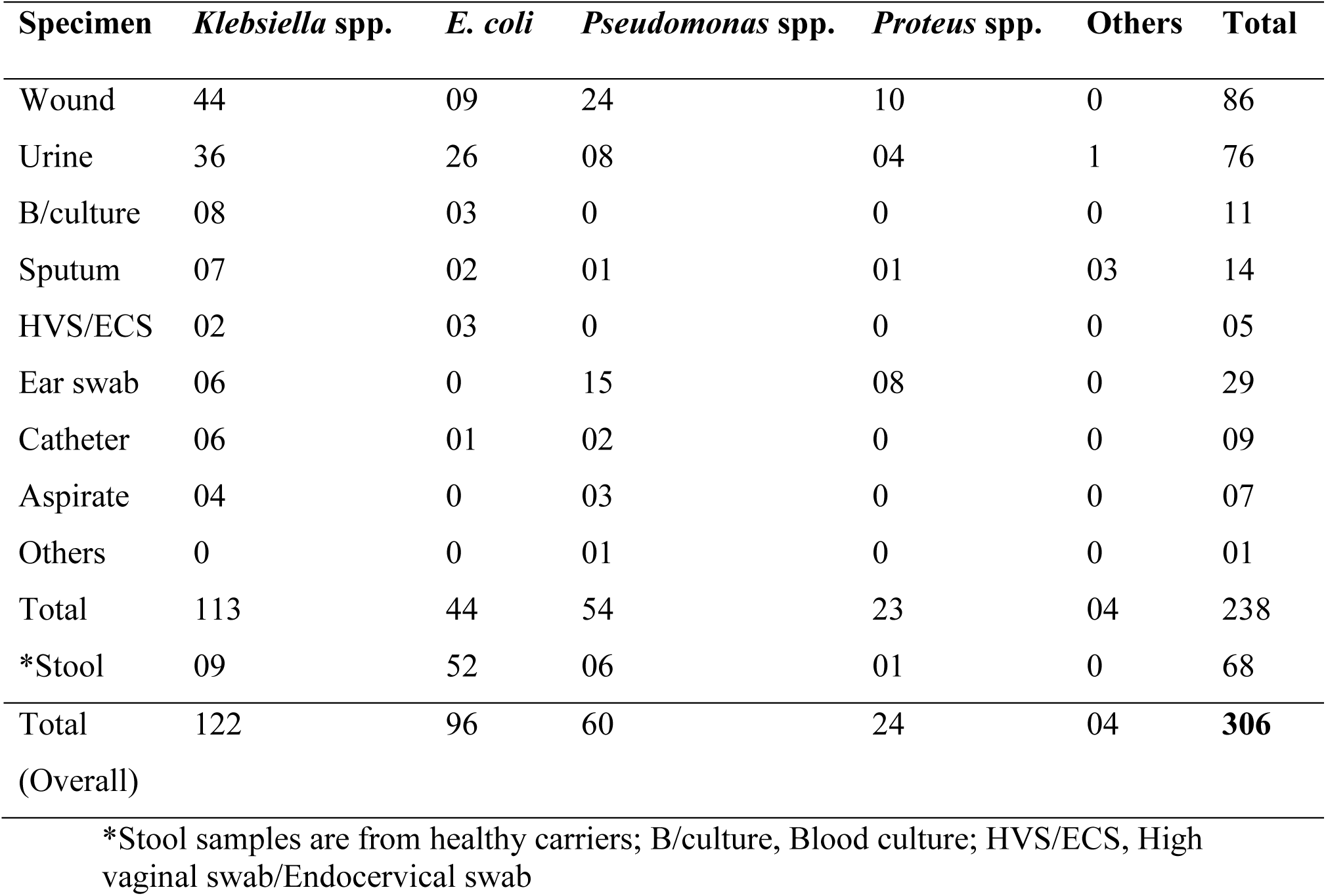
Distribution and sources of bacterial isolates.

**Figure 1.**
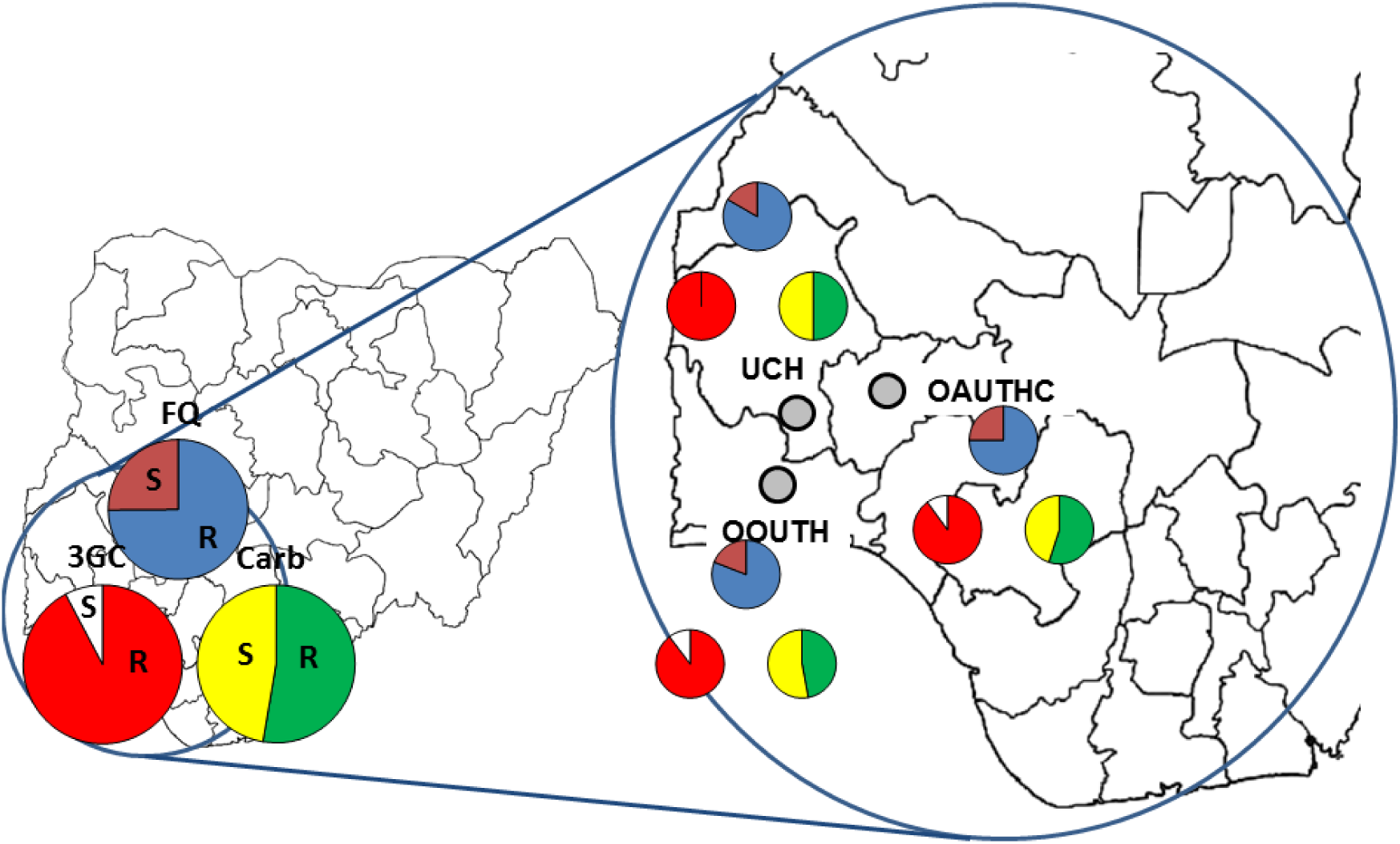
Pie charts show the proportion of isolates classified as resistant (**R**) or sensitive (**S**) to fluoroquinolones (FQ, blue and red), third generation cephalosporins (3GC, red and white) and carbapenems (Carb, green and yellow). The left hand of the figure shows data from the whole study (including all species and both years). The expanded insert (right side) shows data from each of the three hospitals (grey circles).

In addition, isolates from stool samples sent for routine examination from patients but without infection were also collected. Isolates were non-duplicate and unbiased (i.e. no selective criteria beyond being a Gram negative bacterial species were applied) and were randomly collected from the hospital laboratories in 2011 and 2013. A total of 306 isolates were retained and no information about the antibiotic susceptibility of any isolates was used as an inclusion criterion. The isolates comprised *E. coli, Klebsiella* spp*, Pseudomonas aeruginosa, Proteus* spp and others (*Serratia* and *Citrobacter* spp,). Species assignments were confirmed for all isolates using standard biochemical tests and API 20E strips (BioMérieux, Basingstoke, UK) for Enterobacteriaceae.

### Antibiotic susceptibility testing

Susceptibility of all isolates to a panel of antibiotic classes commonly used in these hospitals such as fluoroquinolones, third generation cephalosporins and carbapenem were determined by the disk diffusion method on Mueller–Hinton agar using antibiotic disks from Oxoid Ltd. (Basingstoke, UK) according to the recommendations of EUCAST and interpreted according to EUCAST breakpoints[17]. All susceptibility testing experiments included the control organisms *E. coli* NCTC 10418 and *P. aeruginosa* NCTC 10662.

### Identification of carbapenemase production

The Enterobacteriaceae isolates were tested for production of a carbapenemase using the disc based ‘Carbapenemase detection set’ from Mast Group (Bootle, UK) and interpreted using the ‘carbapenemase detection set calculator’ tool as per the manufacturers guidelines.

### Identification of known carbapenemase genes

PCR and sequencing were used to identify genes encoding various known beta-lactamases (including carbapenemases, NDM, VIM, KPC and GES). Primers used are shown in Table S1 having previously extracted DNA by crude boiling method.

### RAPD PCR

A random amplified polymorphic DNA typing approach was used for each species as a rapid and inexpensive way to assess the diversity of strains within each population. Primers and conditions are given in Table S1.

### Whole genome sequencing and bioinformatics

To characterise the strain types, plasmid content and nature of drug resistance genes present in the collection, 10 isolates (due to paucity of fund) were selected for whole genome sequencing based on their antimicrobial profiling, carbapenemase production, genotypes, clinical specimen and source and species. DNA was extracted with the QIAamp DNA Mini Kit according to manufacturer instruction. Paired-end Illumina sequencing was used to generate high-quality 250 bp reads. Assembly used Velvet [18] and contigs were re-ordered against relevant reference genomes using MAUVE[19]. Assemblies were annotated using RAST (http://rast.nmpdr.org/rast.cgi). Assemblies were used to search for plasmid content and to determine MLST types using the ‘PlasmidFinder’ and ‘MLST’ tools hosted at the Centre for Genomic Epidemiology (https://cge.cbs.dtu.dk/services/PlasmidFinder and http://cge.cbs.dtu.dk/services/MLST). The ‘Comprehensive Antibiotic Resistance Database’, CARD was searched to identify antibiotic resistance genes[20]. Specific assembly of plasmids was attempted using ‘plasmidSPAdes’ and plasmid content identified by plasmid network reconstruction using ‘PLACNET’ (http://castillo.dicom.unican.es/request/)[21]. When necessary, reads were mapped against assemblies using Bowtie [22] and visualized in Artemis[23].

## RESULTS

### Antimicrobial resistance in the 307 Nigerian isolates

The percentage of the entire collection of 306 isolates that were resistant to fluoroquinolones, third generation cephalosporins and carbapenems being 78.1%, 92.2% and 52.3%, respectively (Tables 2 and 3). This pattern of high numbers of clinical isolates being resistant to these classes of drug was very similar in all three study sites (fluoroquinolone resistance was seen in 75 - 83% of isolates, cephalosporin resistance in 90 - 100% of isolates and carbapenem resistance in 50 - 55% of isolates). Of concern was the observation that the percentage of isolates from stool of uninfected patients, i.e. those being carried as commensal bacteria that were resistant to third generation cephalosporins (100%) and carbapenems (69.1%) was higher than in isolates from patients being treated for an infection. This suggests that extremely high numbers of antibiotic resistant bacteria are prevalent in the community and that multidrug resistance is not restricted to isolates found in the hospital environment.

**Table 2.** Summary of antimicrobial disk susceptibility testing of 306 bacterial isolates.

| Organism, n | FQ |  | 3GS |  | CARB |  |
| --- | --- | --- | --- | --- | --- | --- |
|  | S | R | S | R | S | R |
| *K. pneumoniae*, 122 | 30 (24.6) | 92 (75.4) | 09 (7.4) | 113 (92.6) | 60 (49.2) | 62 (50.8) |
| *E. coli*, 96, | 18 (18.6) | 78 (81.4) | 06 (6.3) | 90 (93.7) | 39 (40.6) | 57 (59.4) |
| *Pseudomonas* spp., 60 | 14 (23.3) | 46 (76.7) | 5 (8.3) | 55 (91.7) | 29 (48.3) | 31 (51.7) |
| *Proteus* spp., 24 | 4 (16.7) | 20 (83.3) | 3 (12.5) | 21 (87.5) | 13 (54.2) | 11 (45.8) |
| Others, 04 | 01 (25.0) | 03 (75.0) | 01 (25.0) | 03 (75.0) | 04 (100) | 0 (0) |

FQ, fluoroquinolone; 3GC, third generation cephalosporin; CARB, carbapenem; (), number in parentheses are percentages

**Table 3.** Overview of susceptibility testing to clinically important antibiotics.

|  | Number of isolates | Percentage of isolates resistant |  |  |
| --- | --- | --- | --- | --- |
|  |  | FQ | 3GC | CARB |
| Total isolates | 306 | 78.1 | 92.2 | 52.6 |
| Clinical samples | 238 | 80.4 | 90 | 47.8 |
| Stool (healthy individuals) | 68 | 69.1 | 100 | 69.1 |
| *Klebsiella* spp | 122 | 75.4 | 92.6 | 50.8 |
| *E. coli* | 96 | 81.4 | 93.7 | 59.4 |
| *Pseudomonas* spp | 60 | 76.7 | 91.7 | 51.7 |
| *Proteus* spp | 24 | 83.3 | 87.5 | 45.8 |
| Others | 4 | 75.0 | 75.0 | 0 |
| 2011 | 182 | 78.5 | 96.6 | 59.3 |
| 2013 | 124 | 76.9 | 86.8 | 43.0 |
| Hospital I (UCH) | 212 | 80.9 | 100 | 50 |
| Hospital II (OOUTH) | 6 | 83.3 | 100 | 50 |
| Hospital III (OAUTHC) | 20 | 75 | 90 | 55 |

FQ, fluoroquinolone; 3GC, third generation cephalosporin; CARB, carbapene

When stratified by species, *E. coli* were most commonly resistant to third generation cephalosporins (93.7%) and carbapenems (59.4%), followed by *Pseudomonas* species (where 91.7% of isolates were resistant to third generation cephalosporins and 51.7% of isolates were resistant to carbapenems) (Tables 2, 3 and 4). Of the 307 isolates, no species had 50% or more of isolates which were sensitive to all three classes of antibiotic. The isolates of *Proteus* (n=24) were least likely to be carbapenem resistant; nonetheless 48% of isolates were resistant to this class of drug.

The isolates in this study were collected in two different years, the percent of fluoroquinolone resistant isolates in 2011 and 2013 was very similar. However, between the two years the percentage of isolates resistant to cephalosporins fell from 97% to 87% and the percentage of carbapenem resistant isolates fell from 59% to 43% (Table 1).

### Typing of isolates

RAPD PCR was used to type 54 of 306 isolates representing 18 each of *E. coli*, *K. pneumoniae* and *P. aeruginosa.* A wide variety of strains were present for each species with only a small number of repeated patterns observed e.g. Figure 2 where 13 RAPD patterns were identified from 18 isolates of *E. coli* demonstrating a lack of dominance by specific clones.

**Figure 2.**
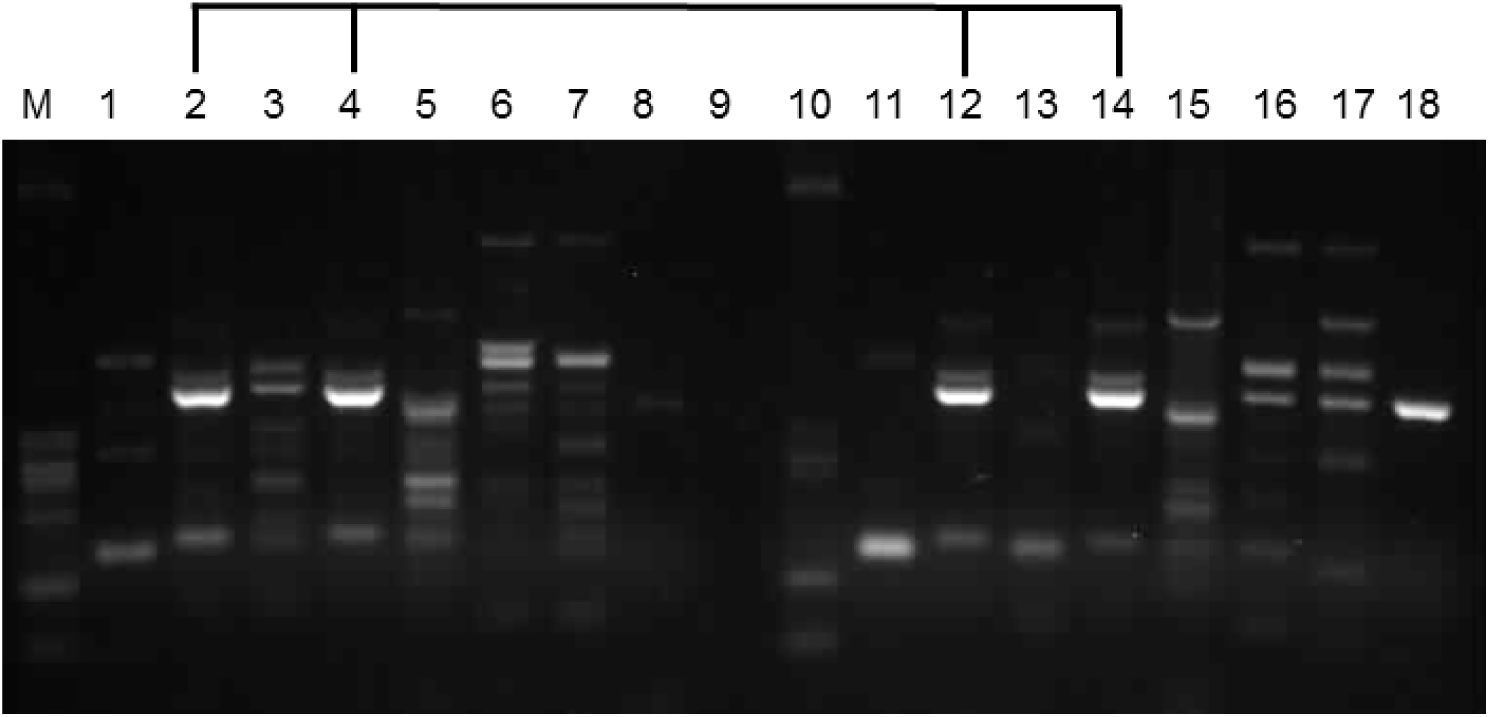
RAPD analysis of a selection of isolates of *E. coli*. The bold line above the gel shows isolates with an identical banding pattern.

### Carbapenemase production and identification of carbapenemase genes

As there was a high prevalence of carbapenem resistance in the isolates, the *E. coli* and *Klebsiella* strains were tested for production of carbapenemases using the ‘carbapenemase detection set’ from Mast Group (Bootle, UK). Only 6% of all isolates produced a carbapenemase (7% of *Klebsiella* and 10% of *E. coli*). Specific carbapenemase genes were amplified by PCR and genes verified by DNA sequencing alleles for all 306 isolates. In agreement with the phenotypic testing, only 19 of the 306 isolates (6.2%) carried a known carbapenemase gene. The PCR revealed the presence of variants of VIM (n=9), GES (n=10) and NDM (n=2) families. These genes were detected in *K. pneumoniae* (n=6), *E. coli* (n=4) and *P. aeruginosa* (n=9) isolates. Two isolates carried two carbapenemase genes (NDM and VIM and GES and VIM). The KPC, IMP or OXA-48 genes were not detected in any isolate.

As 51.3% (157 isolates) of the 306 isolates were resistant to carbapenems but most did not appear to produce a known carbapenemase the presence of other known resistance mechanisms was investigated. The CTX-M genes have been shown to be very common and important in Gram negative isolates among other extended-spectrum β-lactamases around the world, we therefore used primers specific for each of the CTX-M sub-groups were used to detect these genes. Of the 218 *E. coli* and *K. pneumoniae* isolates, 79.4% (173) contained a CTX-M allele; DNA sequencing of a random selection of 40 isolates revealed all to be CTX-M-15. None of these isolates demonstrated de-repression of efflux but all showed either complete loss or reduced production of outer membrane porins (data not shown).

### Characterisation of antibiotic resistance mechanisms and strain types in representative isolates

To investigate the molecular basis of drug resistance ten isolates were chosen for whole genome sequencing. These included representative isolates of the most common species and resistance phenotypes present in the collection. The ten isolates included two *K. pneumoniae*, two *E. coli*, three *P. aeruginosa*, two *P. mirabilis* and one *P. rettgeri* isolate (Table 4). The choice of isolate was informed by susceptibility testing, year of isolation, site of isolation (both geographical and specimen type) and by results of random amplified polymorphic DNA (RAPD) typing. The sequencing identified some globally established strain types in circulation in Nigeria, notably *K. pneumoniae* ST11 and *P. aeruginosa* ST224.

**Table 4.** Strain types, beta-lactamase genes and plasmid replicons present in representative isolates.

| Strain | Species | Sequence<br>Type | MIC (µg/dl) |  |  | Carbapenemase<br>gene | Beta-lactamase genes | Plasmid replicons<br>identified |
| --- | --- | --- | --- | --- | --- | --- | --- | --- |
|  |  |  | IMP | MER | ETP |  |  |  |
| U37 | *K. pneumoniae* | ST11 | 16 | 16 | 64 | NDM-1 | OXA-1, SHV-11, CTX-M-15, | FIB (K), FII (K) |
| U52 | *K. pneumoniae* | ST11 | 16 | 16 | 64 | NDM-1 | OXA-1, SHV-11, CTX-M-15, | FIB (K), FII (K) |
| S46 | *E. coli* | ST226 | 16 | 0.12 | 0.12 | - | AmpC, TEM-1 | FII |
| F124 | *E. coli* | ST156 | 0.5 | 0.06 | 0.06 | - | TEM-1, ACT-7 | Q1 |
| F19 | *P. aeruginosa* | ST244 | 8 | 8 | NA | PDC-1 | - | - |
| F46 | *P. aeruginosa* | ST233 | 32 | 64 | NA | VIM-2, PDC-3 | OXA-33 | - |
| U36 | *P. aeruginosa* | ST233 | 64 | 32 | NA | VIM-2, PDC-3 | OXA-33 | - |
| F10 | *P. mirabilis* | - | 8 | 0.25 | 2 | - | CMY-41, CMY-31, TEM-1 | Q1 |
| F56 | *P. mirabilis* | - | 8 | 0.25 | 0.25 | - | ACT-7, TEM-1 | Q1, Col(BS512) |
| S39 | *P. rettgeri* | - | 16 | 2 | 4 | - | SRT-2 | - |

- Indicates no gene or replicon detected, for sequence type a species where no MLST scheme has been established. N/A;
*Pseudomonas* are intrinsically resistant to ertapenem so no MIC values were determined.

Both the *K. pneumoniae* isolates belonged to ST11 and were from urine of different patients in 2011 from UCH. Both genome assemblies were essentially identical and both carried NDM-1 and CTX-M-15 (Table 4). In addition, both isolates also carried OXA-1 and SHV-11. Interestingly, there was direct evidence for the mobility of the NDM-1 gene in this strain. Whilst the gene encoding NDM-1 was detected by PCR using a boiled colony preparation as a template in both isolates (U52 and U37), in both genome assemblies created after sequencing isolated DNA preparations, it was only initially seen in the genome assembly for U52. Analysis of the genetic location of this gene showed it to be present in a context like that seen by others previously, in an operon with *bleMBL* and associated with *trpF*, *dsbC* and *cutA* (Figure 3). In isolate U37 this region was absent and no sequence reads mapped against the U52 reference (Figure S1) demonstrating likely mobility of this whole region as has been suggested previously [24]. Both IncFIB and FII plasmid replicons were present in both strains supporting a plasmidic context for *bla*NDM-1 (Figure 3). In addition to the beta-lactamase genes, both isolates also carried trimethoprim (*dfrA12*), macrolide (*mphA*) aminoglycoside (*rmtF*) chloramphenicol (*cat*) and sulphonamide (*sul1*) resistance genes. Consistent with fluoroquinolone resistance, mutations in *gyrA* were seen,

**Figure 3.**
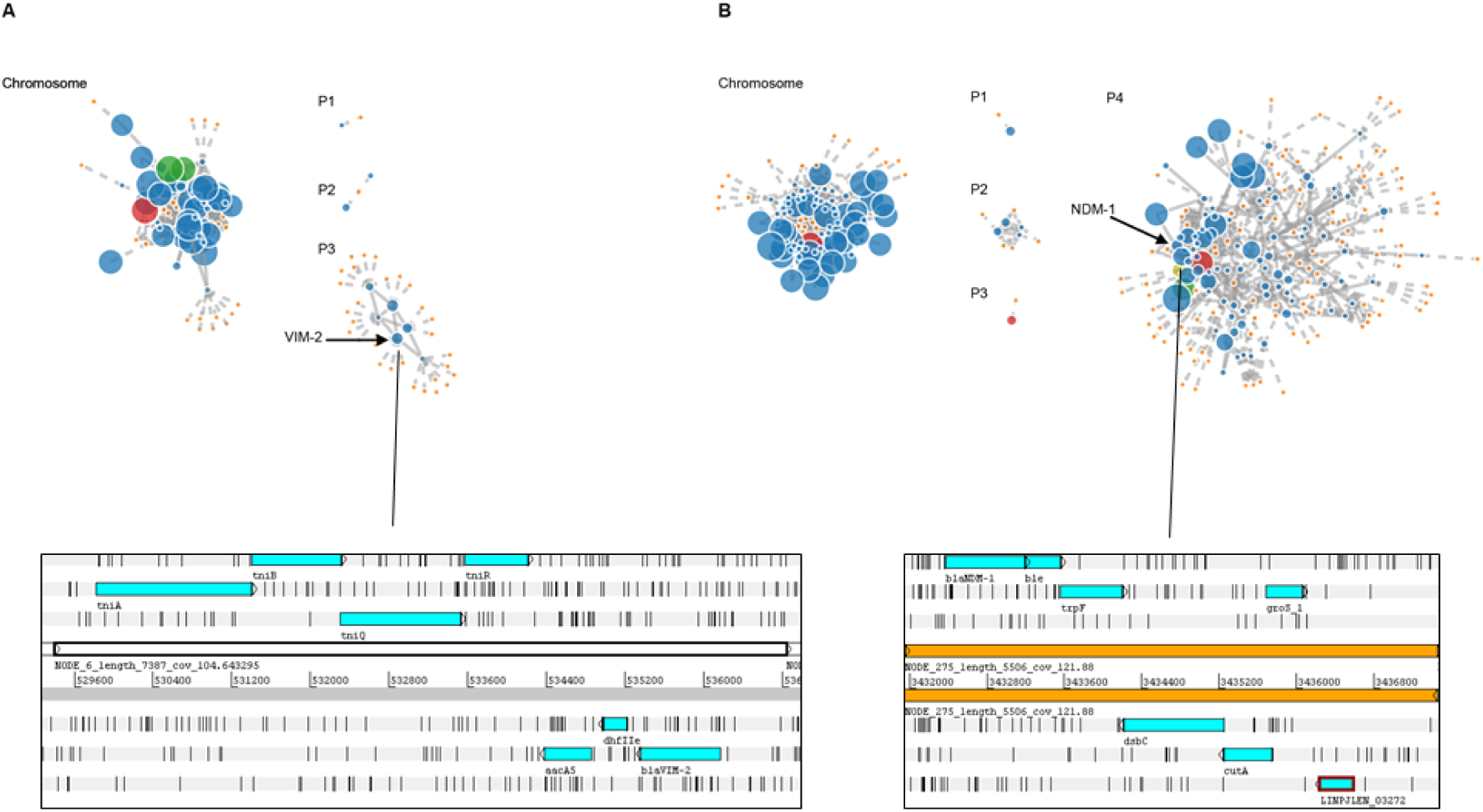
Plasmid content reconstructed using PLACNET. Panel **A** shows network from *P. aeruginosa* F46 with a chromosomal network of 60 contigs totalling ~6.7MB and three discrete plasmid networks; P1 and P2 both <5kb and P3, carrying VIM-2 consisting of 8 contigs of ~31kb. The *bla*VIM-2 gene in *Pseudomonas* was present on a small contig with homology to an integron. Trimethoprim and aminoglycoside resistance genes were also present in the assembled contig. Panel **B** shows network from *K. pneumoniae* U52 with a chromosomal group of 87 contigs totalling 5.3MB and 4 plasmid networks; P1 and P3 both ~5kb, P2 of 9kb and P4 a large network of 90 contigs totalling 1.9 MB which is likely to represent more than one IncF type plasmids which have not been resolved. The *bla*NDM-1 gene in U52 was on a small contig in an operon with the *ble* gene. Immediately downstream were the *trpF*, *dsbC* and *cutA* genes.

One of the *P. aeruginosa* belonged to ST244 and carried the mutant PDC-1 AmpC enzyme as well as genes that contribute to resistance to chloramphenicol (*cmx*, *catB7*), aminoglycosides (*aph(3”)-I1*, *aph(6)-Id*) and fosfomycin (*fosA*). The other two isolates were both members of ST233 and both carried PDC-3. These latter two isolates also carried VIM-2 and OXA-33, were of the same MLST type and both isolated from OOUTH although isolated two years apart. Reads from the F46 strain carrying VIM-2 were assembled using both Velvet and SPAdes (using the plasmidSPAdes option); both resulted in assemblies with the VIM-2 gene present on a contig of ~7000bp. When this sequence was compared with known sequences in Genbank using the BLAST algorithm a perfect match for an integron carrying VIM-2 was found (accession number KT768111.1). Figure 3 shows a plasmid network reconstruction and the genetic context of the VIM-2 genes in these two isolates.

The two *E. coli* strains sequenced belonged to ST226 and ST156. Neither carried known carbapenemase genes although both had multiple mutations within *ampD* suggesting de-repression of the chromosomal *ampC* gene. Both strains also carried TEM-1 and various other mobile resistance genes including genes conferring aminoglycoside resistance (*aph(6)-Id*, *aph(3”)-I1*). An IncFII plasmid replicon was present in isolate S46 (the ST226 isolate).

Two *P. mirabilis* strains were sequenced, isolate F10 carried two CMY genes; CMY-41 reported once previously in a *Citrobacter freundii* isolated from food in Egypt [25] and CMY-31 previously reported in *Klebsiella* and *Salmonella*[26. 27]. A Q1 plasmid replicon was present in F10. This isolate also carried two separate aminoglycoside resistance genes (*aadA5*, *aph(3”)-I1*), as well as chloramphenicol (*catI*), sulphonamide (*sul1*) and plasmidic quinolone resistance genes (*qnrA1*). *P. mirabilis* isolate F56 was found to carry a novel CMY enzyme with a single substitution (of glutamic acid for aspartic acid at codon 144) distinguishing this protein from CMY-48 isolated from *C. freundii*. Isolate F56 also carried a chloramphenicol acetyltansferase gene (*catI*) and three aminoglycoside resistance genes (*aadA5*, *aph(3”)-I1* and *aph(6)-Id*).

The *Providencia* isolate (S39) sequenced carried an SRT-2 AmpC beta-lactamase variant; this has previously been described in *Serratia marcescens* [28]. No other beta-lactamase genes or plasmid replicons were detected in this isolate.

## DISCUSSION

This study suggests that there is a very high prevalence of antibiotic resistance in Nigerian isolates of Gram-negative bacteria to three key classes of antibiotic. A high frequency of resistance to fluoroquinolones and cephalosporins have been seen in other areas of the world increasing the reliance on carbapenems for the treatment of infections caused by Gram negative bacteria. In this study over 50% of the Nigerian isolates in our collection were carbapenem-resistant; empiric use of these antibiotics for the treatment of serious infections is unlikely to be effective. Resistant isolates appear to be widely spread in the community and were not restricted to hospital patients. Isolates from stools of healthy individuals were more likely to be resistant to all three classes of antibiotic tested than those from clinical samples suggesting that the wider Nigerian population commonly carry resistant isolates including carbapenem resistant isolates at a high frequency. From our data, resistance to major antibiotics would appear to be the norm in Gram negative bacteria carried in the Nigerian population.

Characterisation of the mechanisms of carbapenem resistance in our collection of isolates showed that some well-known and globally disseminated carbapenemase genes are in circulation within Nigeria. These included NDM, VIM and GES enzymes. However, less than 10% of the isolates in the study carried a known carbapenemase (according to both molecular and phenotypic testing) and none carried KPC or OXA family carbapenemases. A recent report from Edo state, (in south Nigeria, further east from the locations in this study) has reported the existence of OXA family carbapenemases of OXA-48 and OXA-181 and NDM-1[29]. Carbapenem antibiotics are available in Nigeria but have historically not been widely used in hospital medicine as they have not been part of the common antibiotic formulary. In most Nigerian hospitals third generation cephalosporins, aminoglycosides and fluoroquinolones are the most prescribed antibiotics. There was essentially pan-resistance to the cephalosporins and fluoroquinolones in the isolates. The high level of phenotypic resistance to carbapenems in this collection could be caused by the carriage of currently carbapenemases that were not detectable by the methods used. However, we hypothesize that the very high frequency of carriage of ESBLs (~80% of isolates of Enterobacteriaceae contained CTX-M variants) and AmpC variants in combination with porin loss (in *Pseudomonas* isolates) selected by prolonged and heavy cephalosporin usage are the cause of carbapenem resistance in these isolates. A recent study described Enterobacteriaceae isolates from the USA which were carbapenem resistant without carriage of known carbapenemases) [30](30).

This study covered the South West of Nigeria, where the population density of the country is highest with approximately 50 million people. The study area included major population centers close to other major cities with a diverse population in terms of culture, race, religion and social standing. The major international transportation hubs of Nigeria are also in the South West of the country and over 15,000 international flights leave annually to over 30 different countries and over 8 million passengers fly through Nigeria annually [31]. International destinations are varied with Europe and the Middle East being most common followed by destinations in Asia with a smaller number of flights departing to North and South America [31].

Whilst local antibiotic use is likely to have made an impact on the incidence of antibiotic resistance in the collection of isolates, globally disseminated strain types and resistance genes were identified. This is highly relevant given the mobility of the Nigerian population and the implications for this mobility in global transfer of strains and genes. For example, a recent case report documents a Canadian visitor who suffered a lower leg fracture requiring surgical repair in Nigeria and was repatriated two months later with a wound infected with *Klebsiella*, *Pseudomonas* and *E. coli* isolates all carrying carbapenemases[32].

*K. pneumoniae* strains belonging to ST11 were detected; these have been associated with the carriage of CTX-M-15 and KPC enzymes, mainly in China. ST11 is also a single locus variant from ST258 which has been associated with the international dissemination of KPC enzymes on the pKPQIL plasmid[15]. ST258 isolates have also been recently associated with NDM carriage in India, Sweden and the United Kingdom[33]. In this study, *E. coli* ST226 was recovered from an uninfected patient; this strain type has been found circulating in highly resistant diarrhoeagenic *E. coli* in China. The other *E. coli* ST identified, ST156, has previously been found in Bangladesh[34], and in NDM-1 carrying clinical isolates of *E. coli* from the UK[35]. *P. aeruginosa* clone ST233 has been described as a dominant international ‘high-risk clone’ amongst metallo-β-lactamase-producing *P. aeruginosa* and two VIM-2 positive isolates were found in patients in this study. Isolates of this sequence type have been seen in the UK[16], and have also been reported previously as carrying VIM-2 or IMP-1 in Norway (in an isolate thought to be imported from Ghana)[36], Japan[37], and South Africa[38]. The other *P. aeruginosa* sequence type identified in this study (ST244) is a globally disseminated *P. aeruginosa* clone identified in several countries, including Poland[39], Brazil[40], Spain[41], South Korea[42], the Czech Republic[43], Greece[44], Russia[45], China [46] and Tanzania[47]. ST244 isolates have been found to carry various carbapenemases including IMP and VIM enzymes as well as extended-spectrum ß-lactamases, such as PER-1 and VEB-1[39, 48].

This study demonstrates that antibiotic resistance in Gram-negative bacteria in Nigeria is common place and compromises the effectiveness of the mainstays of broad spectrum empirical therapy. Perhaps most worryingly, this does not appear to be a problem restricted to hospital patients with resistance rates in commensal isolates being carried commensally equally high. The establishment of a reservoir of resistant strains and resistance genes has occurred in Nigeria and this reservoir is likely to be mobilised globally. Our data underpin the urgent requirements for enhanced surveillance of drug-resistance in sub-Saharan Africa and the need for interventions to minimise the selection and transmission of antibiotic resistant Gram-negative bacteria.

## Acknowledgements

We thank the staff within the laboratories of the study site hospitals for help in collection of the isolates used in this study.

## Biographical Sketch

Dr David Ogbolu was born in Ibadan, Nigeria, Africa’s second largest city. After graduating and working as a Biomedical Scientist in 1997 with a specialism in Medical Microbiology he completed a masters degree and subsequently PhD studying mechanisms of antibiotic resistance in Nigerian bacteria. In 2011, David was awarded a Newton Fellowship from the Royal Society to continue his studies and further collaborations with colleagues in the UK. David is now a Senior Lecturer at Ladoke Akintola University of Technology.

## Figure Legends

**Table (S1).**
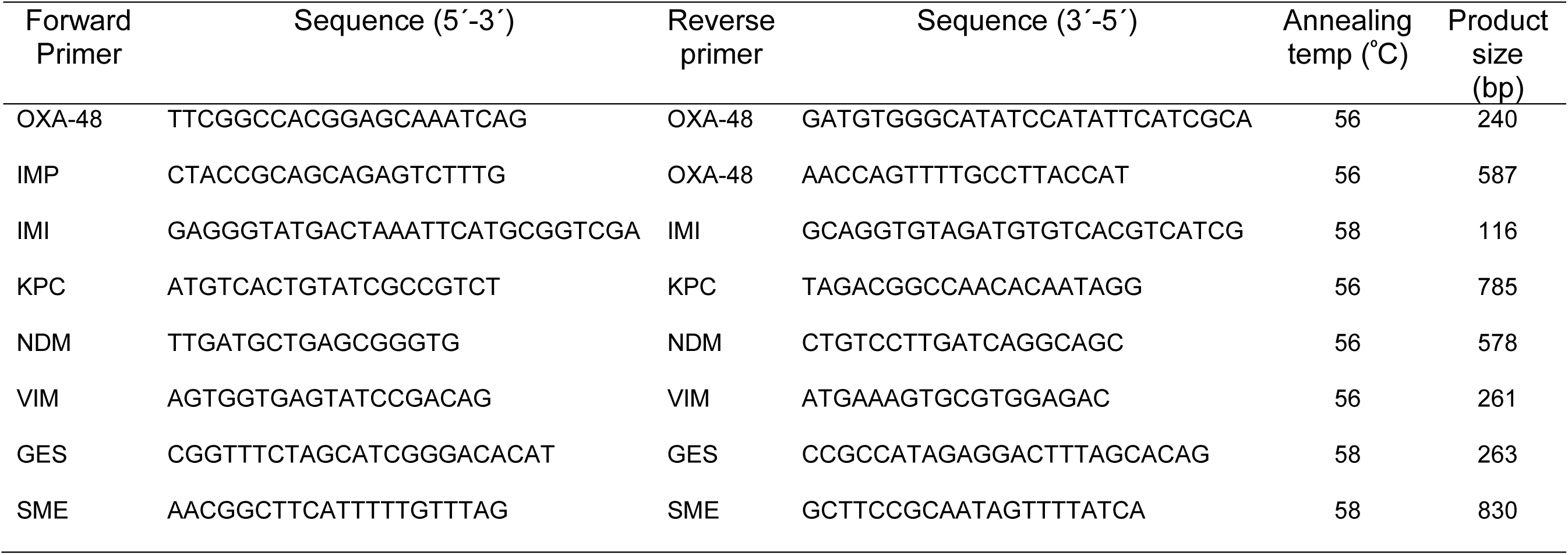
Primers used in this study for the amplification of carbapenemase genes.

## References

1. Akinkunmi EO, Adesunkanmi A-R, Lamikanra A. Pattern of pathogens from surgical wound infections in a Nigerian hospital and their antimicrobial susceptibility profiles. Afr Health Sci. 2014 Dec; 14(4):802–9.

2. Nwankwo EO, Mofolorunsho CK, Akande AO. Aetiological agents of surgical site infection in a specialist hospital in Kano, north-western Nigeria. Tanzan J Health Res. 2014 Oct; 16(4):289–95.

3. Ogbolu DO, Daini O A, Ogunledun a, Alli a O, Webber M a. High levels of multidrug resistance in clinical isolates of Gram-negative pathogens from Nigeria. Int J Antimicrob Agents. Elsevier B.V.; 2011 Jan;37(1):62–6.

4. Ogbolu DO, Webber MA. High-level and novel mechanisms of carbapenem resistance in Gram-negative bacteria from tertiary hospitals in Nigeria. Int J Antimicrob Agents. 2014 May;43(5):412–7.

5. O’Neill J. Tackling drug-resistant infections globally: final report and recommendations the review on antimicrobial resistance. 2016; doi. 10.1016/j.jpha.2015.11.005

6. Blair JMA, Webber MA, Baylay AJ, Ogbolu DO, Piddock LJV. Molecular mechanisms of antibiotic resistance. Nat Rev Microbiol. Nature Publishing Group; 2011;47(14):4055–61.

7. Mshana SE, Matee M, Rweyemamu M. Antimicrobial resistance in human and animal pathogens in Zambia, Democratic Republic of Congo, Mozambique and Tanzania: an urgent need of a sustainable surveillance system. Ann Clin Microbiol Antimicrob. 2013;12:28.

8. Aibinu IE, Ohaegbulam VC, Adenipekun EA, Ogunsola FT, Odugbemi TO, Mee BJ. Extended-spectrum beta-lactamase enzymes in clinical isolates of Enterobacter species from Lagos, Nigeria. J Clin Microbiol. 2003 May;41(5):2197–200.

9. Holt KE, Wertheim H, Zadoks RN, Baker S, Whitehouse CA, Dance D, et al. Genomic analysis of diversity, population structure, virulence, and antimicrobial resistance in Klebsiella pneumoniae, an urgent threat to public health. Proc Natl Acad Sci U S A. 2015 Jul 7;112(27):E3574-81.

10. Adriaenssens N, Coenen S, Versporten A, Muller A, Vankerckhoven V, Goossens H, et al. European Surveillance of Antimicrobial Consumption (ESAC): quality appraisal of antibiotic use in Europe. J Antimicrob Chemother. Oxford University Press; 2011 Dec;66 Suppl 6(suppl 6):vi71-77.

11. Miliani K, L’Hériteau F, Lacavé L, Carbonne A, Astagneau P, Antimicrobial Surveillance Network Study Group. Imipenem and ciprofloxacin consumption as factors associated with high incidence rates of resistant Pseudomonas aeruginosa in hospitals in northern France. J Hosp Infect. 2011 Apr;77(4):343–7.

12. Pournaras S, Koumaki V, Spanakis N, Gennimata V, Tsakris A. Current perspectives on tigecycline resistance in Enterobacteriaceae: susceptibility testing issues and mechanisms of resistance. Int J Antimicrob Agents. 2016 Jul;48(1):11–8.

13. Osei Sekyere J, Govinden U, Bester LA, Essack SY. Colistin and tigecycline resistance in carbapenemase-producing Gram-negative bacteria: emerging resistance mechanisms and detection methods. J Appl Microbiol. 2016 May 7;

14. Zmarlicka MT, Nailor MD, Nicolau DP. Impact of the New Delhi metallo-beta-lactamase on beta-lactam antibiotics. Infect Drug Resist. 2015;8:297–309.

15. Findlay J, Hopkins KL, Doumith M, Meunier D, Wiuff C, Hill R, et al. KPC enzymes in the UK: an analysis of the first 160 cases outside the North-West region. J Antimicrob Chemother. 2016 May;71(5):1199–206.

16. Wright LL, Turton JF, Livermore DM, Hopkins KL, Woodford N. Dominance of international “high-risk clones” among metallo-β-lactamase-producing Pseudomonas aeruginosa in the UK. J Antimicrob Chemother. 2015 Jan;70(1):103–10.

17. European Committee on Antimicrobial Susceptibility Testing Breakpoint tables for interpretation of MICs and zone diameters. Accessed from: http://www.eucast.org/fileadmin/src/media/PDFs/EUCAST_files/Breakpoint_tables/v5.0_Breakpoint_Table_01.pdf

18. Zerbino DR, Birney E. Velvet: algorithms for de novo short read assembly using de Bruijn graphs. Genome Res. 2008 May; 18(5):821–9.

19. Rissman AI, Mau B, Biehl BS, Darling AE, Glasner JD, Perna NT. Reordering contigs of draft genomes using the Mauve aligner. Bioinformatics. 2009 Aug 15;25(16):2071–3.

20. McArthur AG, Waglechner N, Nizam F, Yan A, Azad MA, Baylay AJ, et al. The comprehensive antibiotic resistance database. Antimicrob Agents Chemother. 2013 Jul;57(7):3348–57.

21. Antipov D, Hartwick N, Shen M, Raiko M, Lapidus A, Pevzner PA. plasmidSPAdes: Assembling Plasmids from Whole Genome Sequencing Data. Bioinformatics. 2016 Jul 27;

22. Langmead B, Trapnell C, Pop M, Salzberg SL. Ultrafast and memory-efficient alignment of short DNA sequences to the human genome. Genome Biol. 2009;10(3):R25.

23. Rutherford K, Parkhill J, Crook J, Horsnell T, Rice P, Rajandream MA, et al. Artemis: sequence visualization and annotation. Bioinformatics. 2000 Oct; 16(10):944–5.

24. Dortet L, Nordmann P, Poirel L. Association of the emerging carbapenemase NDM-1 with a bleomycin resistance protein in Enterobacteriaceae and Acinetobacter baumannii. Antimicrob Agents Chemother. 2012 Apr;56(4):1693–7.

25. Hammad AM, Ishida Y, Shimamoto T. Prevalence and molecular characterization of ampicillin-resistant Enterobacteriaceae isolated from traditional Egyptian Domiati cheese. J Food Prot. 2009 Mar;72(3):624–30.

26. Zioga A, Whichard JM, Kotsakis SD, Tzouvelekis LS, Tzelepi E, Miriagou V. CMY-31 and CMY-36 cephalosporinases encoded by ColE1-like plasmids. Antimicrob Agents Chemother. 2009 Mar;53(3):1256–9.

27. Tsakris A, Poulou A, Markou F, Pitiriga V, Piperaki E-T, Kristo I, et al. Dissemination of clinical isolates of Klebsiella oxytoca harboring CMY-31, VIM-1, and a New OXY-2-type variant in the community. Antimicrob Agents Chemother. 2011 Jul;55(7):3164–8.

28. Wu L-T, Tsou M-F, Wu H-J, Chen H-E, Chuang Y-C, Yu W-L. Survey of CTXM-3 extended-spectrum β-lactamase (ESBL) among cefotaxime-resistant Serratia marcescens at a medical center in middle Taiwan. Diagn Microbiol Infect Dis. 2004;49(2):125–9.

29. Jesumirhewe C, Springer B, Lepuschitz S, Allerberger F, Ruppitsch W. Carbapenemase-producing Enterobacteriaceae-isolates from Edo State, Nigeria. Antimicrob Agents Chemother. American Society for Microbiology; 2017 Aug 12;61(8):AAC.00255-17.

30. Cerqueira GC, Earl AM, Ernst CM, Grad YH, Dekker JP, Feldgarden M, et al. Multi-institute analysis of carbapenem resistance reveals remarkable diversity, unexplained mechanisms, and limited clonal outbreaks. Proc Natl Acad Sci. 2017 Jan 31;114(5):1135–40.

31. Economic Benefits from Air Transport in Nigeria Nigeria country report.

32. Walkty A, Gilmour M, Simner P, Embil JM, Boyd D, Mulvey M, et al. Isolation of multiple carbapenemase-producing Gram-negative bacilli from a patient recently hospitalized in Nigeria. Diagn Microbiol Infect Dis. 2015 Apr;81(4):296–8.

33. Giske CG, Fröding I, Hasan CM, Turlej-Rogacka A, Toleman M, Livermore D, et al. Diverse sequence types of Klebsiella pneumoniae contribute to the dissemination of blaNDM-1 in India, Sweden, and the United Kingdom. Antimicrob Agents Chemother. 2012 May;56(5):2735–8.

34. Rashid M, Rakib MM, Hasan B. Antimicrobial-resistant and ESBL-producing Escherichia coli in different ecological niches in Bangladesh. Infect Ecol Epidemiol. 2015;5:26712.

35. Mushtaq S, Irfan S, Sarma JB, Doumith M, Pike R, Pitout J, et al. Phylogenetic diversity of Escherichia coli strains producing NDM-type carbapenemases. J Antimicrob Chemother. 2011 Sep;66(9):2002–5.

36. Samuelsen O, Toleman MA, Sundsfjord A, Rydberg J, Leegaard TM, Walder M, et al. Molecular epidemiology of metallo-beta-lactamase-producing Pseudomonas aeruginosa isolates from Norway and Sweden shows import of international clones and local clonal expansion. Antimicrob Agents Chemother. 2010 Jan;54(1):346–52.

37. Tsutsui A, Suzuki S, Yamane K, Matsui M, Konda T, Marui E, et al. Genotypes and infection sites in an outbreak of multidrug-resistant Pseudomonas aeruginosa. J Hosp Infect. 2011 Aug;78(4):317–22.

38. Koh TH, Khoo CT, Tan TT, Arshad MABM, Ang LP, Lau LJ, et al. Multilocus sequence types of carbapenem-resistant Pseudomonas aeruginosa in Singapore carrying metallo-beta-lactamase genes, including the novel bla(IMP-26) gene. J Clin Microbiol. 2010 Jul;48(7):2563–4.

39. Empel J, Filczak K, Mrówka A, Hryniewicz W, Livermore DM, Gniadkowski M. Outbreak of Pseudomonas aeruginosa infections with PER-1 extended-spectrum beta-lactamase in Warsaw, Poland: further evidence for an international clonal complex. J Clin Microbiol. 2007 Sep;45(9):2829–34.

40. Fonseca EL da, Freitas F dos S, Vicente ACP. The colistin-only-sensitive Brazilian Pseudomonas aeruginosa clone SP (sequence type 277) is spread worldwide. Antimicrob Agents Chemother. 2010 Jun;54(6):2743.

41. García-Castillo M, Del Campo R, Morosini MI, Riera E, Cabot G, Willems R, et al. Wide dispersion of ST175 clone despite high genetic diversity of carbapenem-nonsusceptible Pseudomonas aeruginosa clinical strains in 16 Spanish hospitals. J Clin Microbiol. 2011 Aug;49(8):2905–10.

42. Lee J-Y, Song J-H, Ko KS. Identification of nonclonal Pseudomonas aeruginosa isolates with reduced colistin susceptibility in Korea. Microb Drug Resist. 2011 Jun;17(2):299–304.

43. Nemec A, Krizova L, Maixnerova M, Musilek M. Multidrug-resistant epidemic clones among bloodstream isolates of Pseudomonas aeruginosa in the Czech Republic. Res Microbiol. 2010 Apr; 161(3):234–42.

44. Liakopoulos A, Mavroidi A, Katsifas EA, Theodosiou A, Karagouni AD, Miriagou V, et al. Carbapenemase-producing Pseudomonas aeruginosa from central Greece: molecular epidemiology and genetic analysis of class I integrons. BMC Infect Dis. 2013;13:505.

45. Edelstein M V, Skleenova EN, Shevchenko O V, D’souza JW, Tapalski D V, Azizov IS, et al. Spread of extensively resistant VIM-2-positive ST235 Pseudomonas aeruginosa in Belarus, Kazakhstan, and Russia: a longitudinal epidemiological and clinical study. Lancet Infect Dis. 2013 Oct; 13(10):867–76.

46. Chen Y, Sun M, Wang M, Lu Y, Yan Z. Dissemination of IMP-6-producing Pseudomonas aeruginosa ST244 in multiple cities in China. Eur J Clin Microbiol Infect Dis. 2014 Jul;33(7):1181–7.

47. Moyo S, Haldorsen B, Aboud S, Blomberg B, Maselle SY, Sundsfjord A, et al. Identification of VIM-2-producing Pseudomonas aeruginosa from Tanzania is associated with sequence types 244 and 640 and the location of blaVIM-2 in a TniC integron. Antimicrob Agents Chemother. 2015 Jan;59(1):682–5.

48. Vatcheva-Dobrevska R, Mulet X, Ivanov I, Zamorano L, Dobreva E, Velinov T, et al. Molecular epidemiology and multidrug resistance mechanisms of Pseudomonas aeruginosa isolates from Bulgarian hospitals. Microb Drug Resist. 2013 Oct; 19(5):355–61.

